# Aspiration-assisted freeform bioprinting of mesenchymal stem cell spheroids within alginate microgels

**DOI:** 10.1101/2021.07.13.452209

**Authors:** Myoung Hwan Kim, Dishary Banerjee, Nazmiye Celik, Ibrahim T Ozbolat

**Affiliations:** Department of Biomedical Engineering, Pennsylvania State University, University Park, PA, USA; The Huck Institutes of the Life Sciences, Pennsylvania State University, University Park, PA, USA; Engineering Science and Mechanics Department, Penn State University, University Park, PA, USA; Materials Research Institute, Pennsylvania State University, University Park, PA, USA; Department of Neurosurgery, Pennsylvania State College of Medicine, Hershey, PA, USA

**Keywords:** bioprinting, aspiration-assisted freeform bioprinting, mesenchymal stem cell spheroids, bone, alginate microgels

## Abstract

Aspiration-assisted freeform bioprinting (AAfB) has emerged as a promising technique for precise placement of tissue spheroids in three-dimensional (3D) space for fabrication of tissues. For successful embedded bioprinting using AAfB, an ideal support bath should possess shear-thinning behavior and yield-stress to obtain tightly fused assembly of bioprinted spheroids. Several studies have demonstrated support baths for embedded bioprinting, but these materials pose major challenges due to their low biocompatibility, opaqueness, complex and prolonged preparation procedures, and limited spheroid fusion efficacy. In this study, to circumvent the aforementioned limitations, we present the feasibility of AAfB of human mesenchymal stem cell (hMSC) spheroids in alginate microgels as a support bath. First, alginate microgels were prepared with different particle sizes modulated by blending time and concentration, followed by determination of the optimal bioprinting conditions by the assessment of rheological properties, bioprintability, and spheroid fusion efficiency. The bioprinted and consequently self-assembled tissue structures made of hMSC spheroids were osteogenically induced for bone tissue formation. Alongside, we investigated the effects of peripheral blood monocyte-derived osteoclast incorporation into the hMSC spheroids in heterotypic bone tissue formation. We demonstrated that alginate microgels enabled unprecedented positional accuracy (~5%), transparency for visualization, and improved fusion efficiency (~97%) of bioprinted hMSC spheroids for bone fabrication. This study demonstrates the feasibility of using alginate microgels as a support bath for many different applications including but not limited to freeform bioprinting of spheroids, cell-laden hydrogels, and fugitive inks to form viable tissue constructs.

## 1 Introduction

Bioprinting is a powerful technology enabling the fabrication of anatomically correct tissue constructs in a precise and rapid manner (1). Despite the great progress in the field of bioprinting over the last decade (2), the utilized bioinks have been mainly limited to cell-laden hydrogels. Although hydrogels possess favorable properties for bioprinting, such as biocompatibility, biodegradability, and shear thinning characteristics, the cell density that bioink can carry is usually limited up to tens of millions per milliliter, which is usually less than physiologically-relevant cell densities (3). As a result of being individually encapsulated and confined in a hydrogel network, bioprinted cells experience limited inter- and intra-cellular interactions (4). Therefore, cell aggregates, such as tissue spheroids, have been considered a promising bioink candidate as they possess adequate measurable properties (such as surface tension, stiffness, etc.) along with physiologically-relevant cell densities (3).

Multiple bioprinting techniques including extrusion-based bioprinting (5), droplet-based bioprinting (6), Kenzan method (7), and aspiration-assisted bioprinting (8) have been explored for bioprinting of cell aggregates like spheroids (3), organoids (5), cell pellet (9), and tissue strands (10). Despite these advances and considerable research, three-dimensional (3D) bioprinting of tissue spheroids in highly intricate freeform shapes has been a long-standing problem. In this regard, we recently introduced the concept of bioprinting spheroids in yield-stress gels, referred to aspiration-assisted freeform bioprinting (AAfB) (11). In this technique, spheroids are picked up by aspiration forces and then transferred into a yield-stress support bath for accurate placement on a target position in 3D. Despite significant progress of using yield-stress support baths in embedded bioprinting of hydrogels like FRESH (Freeform reversible embedding of suspended hydrogels) (12, 13), currently available support formulations pose major limitations for AAfB of spheroids such as limited biocompatibility and spheroid fusion efficiency, opaqueness, cumbersome preparation procedures, and difficulties in removal of assembled spheroids after fusion. For example, in our recent work (11), we utilized commercially available Carbopol and alginate microparticles for AAfB of spheroids, however, Carbopol induced major cell death after three days of culture (~74% of cell viability), dissolved in the cell media quickly, and generated major instability issues during the fusion of spheroids. Whereas alginate microparticles, prepared using standard procedures, demonstrated better spheroid viability but were highly opaque limiting the vision and automation capabilities and the particle size was relatively large limiting the fusion efficiency of spheroids (up to 65%, 65% fusion efficiency indicates the disassembly of 3-4 constructs out of 10).

In this regard, we here present a new preparation procedure for alginate microgels resulting in fine microparticles i) enabling effective transfer of spheroids from the cell media to the gel compartment without inducing any major damage to the spheroid integrity and viability, ii) facilitating a transparent view of the gel domain, which is essential for advanced vision and automation systems, iii) improving the bioprinting accuracy and precision, and iv) providing shear-thinning and self-healing properties for successful transfer and then positioning of spheroids in a highly stable manner ensuring their successful fusion. In this study, we hypothesized that smaller size of Alg microgels would serve as a promising condition for AAfB with improved bioprinting precision and accuracy and enhanced fusion efficiency of tissue spheroids. By modulating the blending time and the concentration of alginate, we achieved ~97% spheroid fusion efficiency and an unprecedented spheroid bioprinting accuracy (~5% with respect to the spheroid size) during freeform bioprinting of spheroids.

Using the optimized alginate microgels, we henceforth demonstrate successful assembly of human mesenchymal stem cell (hMSC) spheroids and their osteogenic differentiation for bone tissue formation. Since osteoclasts also play a significant role in natïve bone remodeling (14, 15), to the best of our knowledge, we also for the very first time demonstrate the incorporation of human peripheral blood monocyte (THP-1) derived osteoclasts into hMSCs to form heterocellular spheroids and investigate their role in self-assembly and consequently in formation of bone tissue.

## 2 Materials and methods

### 2.1 Preparation of alginate (Alg) microgels

To prepare alginate microgels, sodium alginate (Alg, Sigma Aldrich Inc., MO, USA) was dissolved in ultra-purified water at several concentrations (0.5, 1.0, and 2.0% w/v) at room temperature. The homogeneously mixed alginate solution was loaded into a dropping funnel and added dropwise into 4% calcium chloride (CaCl_2_, Sigma Aldrich Inc., MO, USA) solution as a crosslinking agent. After crosslinking of alginate for 30 min, the crosslinked alginate beads were collected, washed thrice with ultra-purified water to remove CaCl_2_ solution and uncrosslinked alginate residues. The alginate beads were blended at 465 × g for 10, 20, and 30 min (B10m, B20m, and B30m) to obtain different particle sizes of Alg microgels using a commercial blender. The resultant microgels were then divided into 50 mL conical tubes and centrifuged at 2000 × g for 5 min. All equipment used for the preparation of Alg microgels were sterilized with 70% ethanol and ultraviolet (UV) light for 30 min.

### 2.2 Morphological evaluation

To visualize the morphology of Alg microgels, Safranin O red staining kit (American MasterTech Scientific, CA, USA) was used according to the manufacturer’s instructions. The stained alginate microgels were placed between a glass microscope slide and cover slide after dilution (1:10 with deionized (DI) water) to clarify microgel particles and imaged using an optical microscope (EVOS FL, Thermo Fisher Scientific, MA, USA). The particle size was investigated by a laser granulometry using a Mastersizer 3000 (Malvern PANalyticals, Worcester, UK) and the volume-weighted mean particle size was quantified. To qualitatively demonstrate the transparency of Alg microgels (B30m) with different concentrations (0.5, 1.0, and 2.0%), each microgels solution was loaded into a 24-well plate positioned on Nittany Lion logos and photographs were taken. The transparency of Alg microgels was also quantitatively assessed through the optical absorbance using a PowerWave X spectrophotometer (BioTek, VT, USA). Alg microgels were loaded into a 96-well plate with a volume of 100 μL for each well and then centrifuged to remove air bubbles. Next, optical absorbance was measured at a fixed wavelength of 560 nm (*n* = 10).

### 2.3 Rheological characterization

Rheological properties of Alg microgels were characterized using a rheometer (MCR 302, Anton Paar, Austria). All rheological measurements were performed in triplicates with a 25 mm parallel-plate geometry at room temperature (22 °C) controlled by a Peltier system. Amplitude sweep test was carried out to evaluate the viscoelasticity of microgels at a constant angular frequency of 10 rad/s in a strain range from 0.01 to 100%. The yield stress of Alg microgels was determined using RheoCompass software (Anton Paar, Austria). The frequency sweep test was carried out in an angular frequency range of 0.1-100 rad/s at a strain in the linear viscoelastic region to examine the storage (G’) and loss modulus (G”) without destroying the sample. To investigate the shear-thinning behavior of Alg microgels, a flow sweep test was achieved in the shear rate range of 0.1-100 s^-1^. A recovery sweep test was performed to evaluate the changes of the viscosity of Alg microgels after applying low and high shear rates at five intervals: (1) a shear rate of 0.1 s^-1^ for 60 sec, (2) a shear rate of 100 s^-1^ for 10 sec, (3) a shear rate of 0.1 s^-1^ for 60 sec, (4) a shear rate of 100 s^-1^ for 10 sec, and (5) a shear rate of 0.1 s^-1^ for 60 sec.

### 2.4 Fabrication and osteogenic differentiation of hMSC spheroids

Human bone marrow-derived mesenchymal stem cells (hMSCs, RoosterBio Inc., MD, USA) were used to fabricate spheroids. Cells at passages from 4 through 7 were used. hMSCs were cultured in mesenchymal stem cell expansion media (R&D Systems, MN, USA) at 37 °C in 5% CO_2_ in a humidified sterile incubator. To prepare hMSC only spheroids, the expanded hMSCs were trypsinized, centrifuged, and transferred to each well of a U-bottom 96-well plate (Greiner Bio One, Austria) with 15,000 cells per well. Spheroids were obtained with a size of 400-600 μm as we reported before (11). Bioprinted spheroids were then cultured in human osteogenic differentiation media (Cell Applications Inc., USA) for 28 days post-bioprinting to enable their differentiation into an osteogenic lineage. The differentiation media was changed every three days.

### 2.5 Fabrication of heterocellular spheroids

Peripheral blood-derived monocytes THP-1 (ATCC, VA, USA) was added to pre-warmed ATCC-formulated RPMI-1640 medium, supplemented with 10% FBS and 0.05 mM 2-mercaptoethanol to reach the desired cell density. After reaching a cell density of ~1 x 10^6^ cells per mL, the suspension media was collected and centrifuged. The cell pellet was resuspended into an osteoclast differentiation media constituting of RPMI 1640 supplemented with 40 ngml^-1^ phorbol 12-myristate 13-acetate (PMA, Abcam, Cambridge, UK), 10 ngml^-1^ receptor activator of nuclear factor-kappa ß ligand (RANKL, Abcam), 10 % FBS, antibiotic/antimycotic, and non-essential amino acids (Gibco, USA) (14–16), mixed with hMSC cells in different ratios of 2:1 and 3:2 hMSCs:THP1 (referred as 2:1 hMSCs:THP1 and 3:2 hMSCs:THP1) and pipetted into each well of a U-bottom 96-well plate to facilitate formation of heterocellular spheroids with ~15,000 cells per well. Cultures were maintained in a mixed media of osteoblast and osteoclast differentiation media at the same ratio as of the cells in heterocellular spheroids at 37 °C under humidified atmosphere of 5% CO_2_. Cell medium was changed every 2–3 days during the entire course of the experiment.

### 2.6 Aspiration-assisted freeform bioprinting

An in-house custom-made bioprinting system, reported before (8, 11, 17), was utilized with modifications for aspiration-assisted freeform bioprinting. As depicted in figure 1(A) and (B), the bioprinting setup was composed of a nozzle, spheroid reservoir, and placement area. In the placement area, a 35 Ø Petri dish was fixed, and a 3D printed pocket was added in the Petri dish to hold Alg microgels during bioprinting. The pocket chamber (15 × 15 × 2 mm^3^) was fabricated by an Ultimaker 3 (Netherlands). The 3D printed pocket was sterilized with 70% ethanol, followed by exposure under UV ray for 1 h. Microgels were loaded into the square pocket before bioprinting. To visualize the bioprinting process in real-time, three microscopic cameras, for top, bottom, and side views, were installed as presented in figure 1(B) and Supplementary videos 1-2. Fabricated spheroids, after 24 h of seeding, were transferred into the reservoir. A straight stainless steel nozzle with a size of 27G (inner diameter: 200 μm, Nordson, OH, USA) was used to aspirate spheroids by applying 70 mmHg aspiration pressure, which was optimized for the preservation of cell viability during bioprinting in our previous work (11). The lifted hMSC spheroids were gently placed in desired positions in microgels (figures 1(A4)-(A6)) at a bioprinting speed of 2.5 mm s^-1^ to generate a dumbbell-shaped structure. The spheroids were then cultured in their expansion media for 2 days to study the effects of microgel properties on bioprinting perfomence properties such as circularity, accuracy, and precision. AAfB was also utilized to bioprint heterocellular spheroids along with hMSC-only spheroids to form triangular tissue patches. Next, the bioprinted hMSC spheroids were cultured in hMSC expansion media. For osteogenically-inducted bioprinted structures, hMSCs only, 2:1 hMSCs:THP1, and 3:2 hMSCs:THP1 groups were cultured in their respective aforementioned differentiation media. The gel compartment was removed after 1 to 5 days of incubation (depending on fusion) by adding 4 w/v% sodium citrate (Sigma Aldrich Inc.) for 15 min as a lyase for alginate (Supplementary figure 5) (18). Bioprinted constructs were washed three times with Dulbecco’s Phosphate Buffered Saline (DPBS, Corning, NY, USA) and cultured with the media that was used for the related tissue structures.

**Figure 1.**
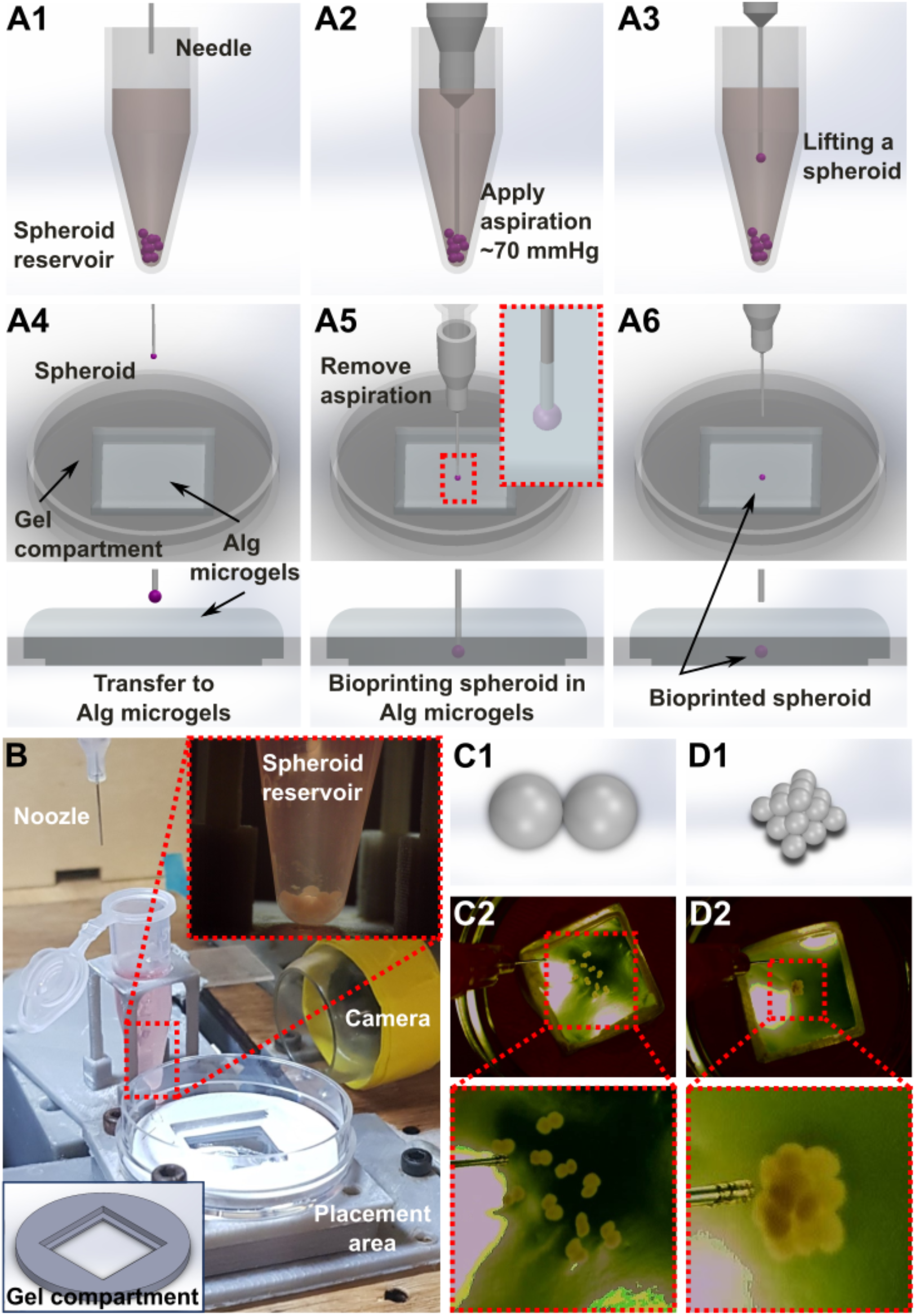
Aspiration-assisted freeform bioprinting process. (A) A schematic demonstrating AAfB, where spheroids were lifted and bioprinted into Alg microgels in an iterative manner. (B) The AAfB setup consisting of a nozzle, spheroid reservoir, cameras, and a gel compartment. (C1, D1) Illustration of dumbbell and pyramid structures, and (C2, D2) bioprinted spheroids for dumbbell and pyramid structures in Alg microgels (0.5% Alg with B30m).

#### 2.6.1 Accuracy and precision measurements

To evaluate the bioprinting accuracy and precision in microgels, hMSC spheroids were bioprinted at a predetermined target position on a micrometer calibration ruler. The calibration ruler was placed at the bottom of a Petri dish and recorded by a USB microscopic camera to monitor the target position. For each microgel group, a total of 10 spheroids were bioprinted and analyzed by ImageJ (National Institutes of Health (NIH), MD, USA), using the following equation:

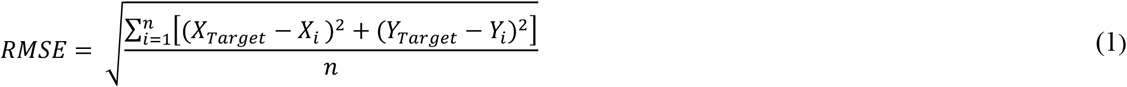

where RMSE (root mean square error) represents the accuracy, *X_Target_* and *Y_Target_* are the *X* and *Y* coordinates of the target position, respectively *X_i_* and *Y_i_* are the position of bioprinted spheroid in *X* and *Y* axes, respectively, and *n* is the sample size. Precision was represented as the square root of the standard deviation. The lower the percentage of accuracy and precision, the higher accuracy and precision.

### 2.7 Biological activities and fusion capability assessment

To evaluate the biocompatibility of Alg microgels, hMSC spheroids were placed into Alg microgels and cultured at 37 °C in 5% CO_2_ in a humidified atmosphere for three days. Cell viability was assessed using LIVE/DEAD staining as we described before (19). Briefly, the cultured spheroids were washed with DPBS, stained with a working solution composed of 2 μM calcein AM (Invitrogen, CA, USA) and 4 μM ethidium homodimer-1 for 30 min in the incubator and then imaged using a Zeiss Axio Observer microscope (Zeiss, Jena, Germany). To observe the fusion dynamics of bioprinted spheroids, pairs of bioprinted hMSC spheroids were monitored using Zeiss Axio Observer microscope for 6 h every 30 min. To quantify the fusion capability, the intersphere angle and contact length between paired spheroids (Supplementary figure 2(A)) were normalized with respect to Day 0 (*n* = 10). The fusion efficiency of bioprinted spheroids was quantified from the number of successfully fused spheroids divided by the total number of bioprinted pairs (*n* = 10) for two days post bioprinting (Supplementary figure 2(B)). To represent the effect of microgel properties on the deformation of bioprinted spheroids, hMSC spheroids (*n* = 9) were bioprinted in each Alg microgels. Spheroids bioprinted in hMSC expansion media without exposure to microgels were used as controls. To quantify the morphology of spheroids, circularity was calculated using ImageJ software (11) using the following equation:

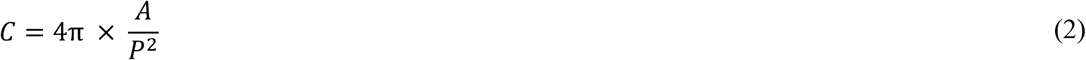

Where *C* represents circularity, *A* is the area, and *P* is the perimeter of the bioprinted spheroid. The value “0” indicated an infinitely elongated polygon and “1” indicated a perfectly circular shape.

### 2.8 Immunohistochemistry and histology of bioprinted spheroids

Bioprinted osteogenically-inducted spheroids were fixed overnight in 4% paraformaldehyde in DPBS (Thermo Scientific), embedded in paraffin (Thermo Scientific). Fixed samples were sectioned into 10-μm thickness using a microtome. Cross-sectioned samples were stained with hematoxylin and eosin (H&E) staining using Leica Autostainer XL (Leica) and then imaged using the Zeiss Axio Observer. Calcium deposition was visualized with a 2% of alizarin red S staining solution. Sectioned samples were stained for 10 min at room temperature. Samples were mounted and imaged using the Zeiss Axio Observer microscope. For immunostaining, bioprinted osteogenically-inducted spheroids were fixed and stained with bone-specific markers. To visualize differentiation of hMSCs and THP1 into osteoblasts and osteoclasts, and cell nuclei, the bioprinted structures were stained with osteoblast differentiation marker Runt-related transcription factor 2 (RUNX2; Abcam), osteoclast differentiation marker tartrate-resistant acid phosphatase (TRAP; Kerafast, MA, USA) and Hoechst (Sigma Aldrich Inc), respectively. The stained spheroids were imaged using the Axio Observer.

### 2.9 Gene expression using quantitative real-time polymerase chain reaction (qRT-PCR)

qRT-PCR was conducted in order to evaluate the osteoblast- and osteoclast-related gene expression profiles in three different groups for bioprinted structures on Day 28. Samples were harvested at Day 28 and homogenized in TRIzol reagent (Life Technologies, CA). Total RNA was isolated from bioprinted structures using PureLink RNA Mini Kit (ThermoFisher) according to the manufacturer’s protocol. RNA concentration was measured using a Nanodrop (Thermo Fisher Scientific, PA). Reverse transcription was performed using AccuPower® CycleScript RT PreMix (BIONEER, Korea) following the manufacturer’s instructions. Gene expression was analyzed quantitatively with SYBR Green (Thermo Fisher Scientific, PA) using a QuantStudio 3 PCR system (Thermo Fisher Scientific). Osteoblast, and osteoclast-related genes tested included OSTERIX (transcription factor Sp7), RUNX2, BSP (Bone sialoprotein), CTSK (Cathepsin K), and MMP9 (Matrix metallopeptidase 9). The reader is referred to Table 1 for the gene sequences. Expression levels for each gene were normalized to glyceraldehyde 3-phosphate dehydrogenase (GAPDH). The fold-change of osteoblast cells at Day 1 were set as 1-fold and values in all groups were normalized with respect to that of the group.

**Table 1.**
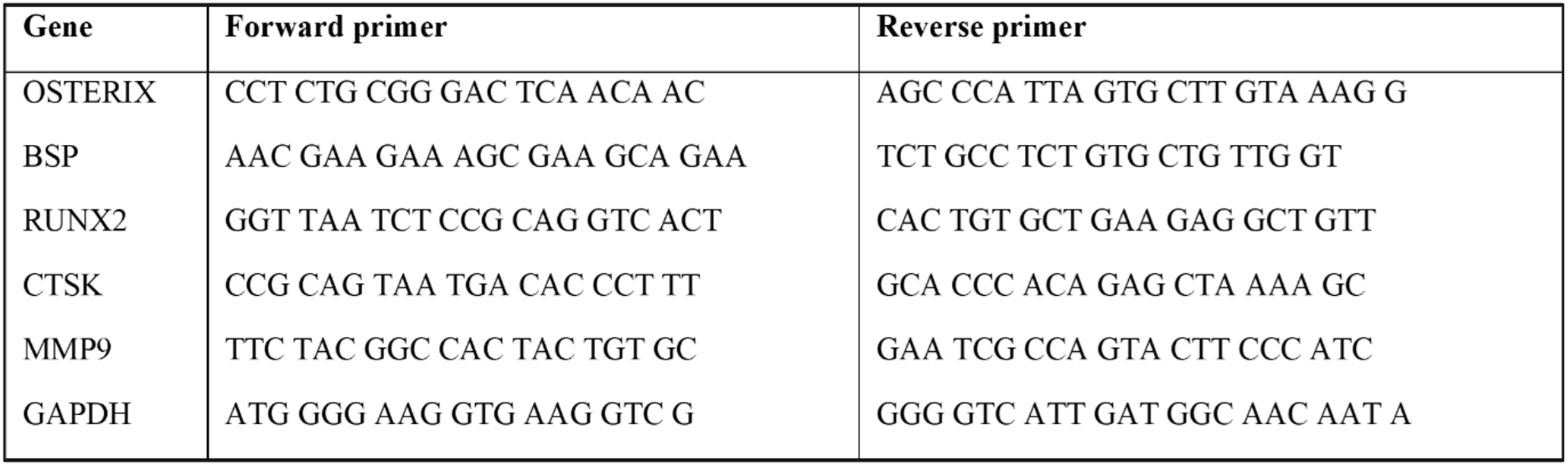
Primers of the measured mRNA for qRT-PCR

### 2.10 Statistical analysis

All data are presented as mean ± standard deviation and analyzed by Minitab 17.3 (Minitab Inc., USA). Multiple comparisons were analyzed by one-way analysis of variance (ANOVA) followed by post-hoc Tukey’s multiple-comparison test to determine the individual differences among the groups. Differences were considered significant at \**p* < 0.05, \*\**p* < 0.01, \*\*\**p* < 0.001. For statistical analysis of bioprinting accuracy and precision, and spheroid fusion efficiency, two-way ANOVA was performed followed by post-hoc Tukey’s multiple-comparison test to determine the individual differences \**p* < 0.05, \*\**p* < 0.01, \*\*\**p* < 0.001.

## 3. Results

### 3.1 Morphological characterization of Alg microgels

To characterize the effect of blending time on the size of Alg microgels, particle diameter and distribution profile of microgels were assessed. Prepared Alg microgels were stained with Safranin O red and observed using optical microscopy (figure 2(A)). The blended Alg microgels consisted of non-spherical particles, mostly in polygonal shapes. The particle size distribution of different Alg concentrations and blending time was analyzed (figure 2(B)). The particle size for all Alg concentrations noticeably decreased as the blending time increased, which was corroborated quantitatively, shown in figure 2(C). For 0.5 and 1.0% Alg microgels, the particle size dropped from ~55 to 31 μm and 121 to 45 μm with increase in blending time, respectively. In contrast, 2.0% Alg had larger particles and it was challenging to reduce the particle size compared to 0.5 and 1.0% Alg. Figure 2(D) shows transparency of Alg microgels (B30m) at different concentrations, where the Nittany Lion logo was blurred more as the concentration increased. In figure 2(E), the optical absorbance demonstrates the transparency of the Alg microgels at a fixed wavelength (560 nm). The absorbance value close to “0” indicated a perfectly transparent case without light scattering. Correlated with figure 2(D), figure 2(E) showed that the absorbance of 0.5 and 1.0% Alg microgels was significantly less than 2.0% Alg microgels, which means that 0.5 and 1.0% Alg microgels were relatively transparent.

**Figure 2.**
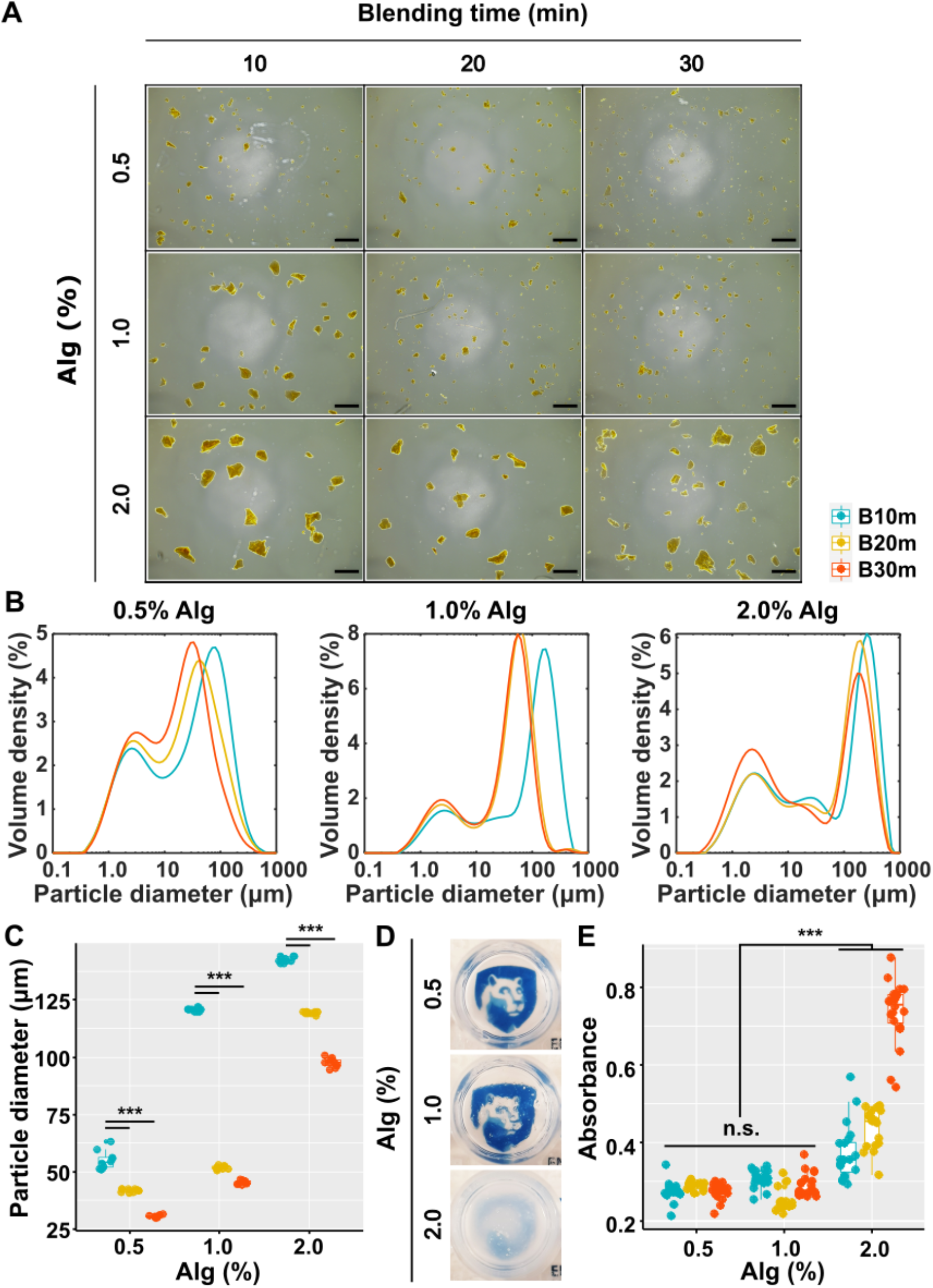
Morphological characterization of Alg microgels at different alginate concentrations and blending time. (A) Microscopic images of stained microgels for visualization (scale bar: 500 μm); (B) the volume density distribution profiles of Alg microgels; (C) Quantified volume-weighted mean particle size (D [4, 3]) (*n* = 10, \*\*\**p* < 0.001); (D) Transparency of Alg microgels (B30m) in a well-plate positioned on Nittany Lion logos, and (E) their optical absorbance (*n* = 10, n.s. not significant, \*\*\**p* < 0.001).

### 3.2 Rheological characterization of Alg microgels

Rheological assessment was carried out to understand the characteristics of Alg microgels at different concentrations (0.5, 1.0, and 2.0%) and blending time (10, 20, and 30 m). To operate AAfB, the supporting bath ideally requires to demonstrate yield-stress, shear-thinning, and self-healing properties. The shear-thinning behavior allows facilitating the needle movement in the bath, enabling precise positioning of bioprinted spheroids in desired positions due to proper yield stress. In parallel, the material recovers from the structural deformities caused by the needle movement owing to its self-healing behavior. The frequency sweep, amplitude sweep, flow sweep, and recovery tests were performed to determine whether Alg microgels exhibit yield-stress, shear-thinning, and self-healing behavior (figure 3).

**Figure 3.**
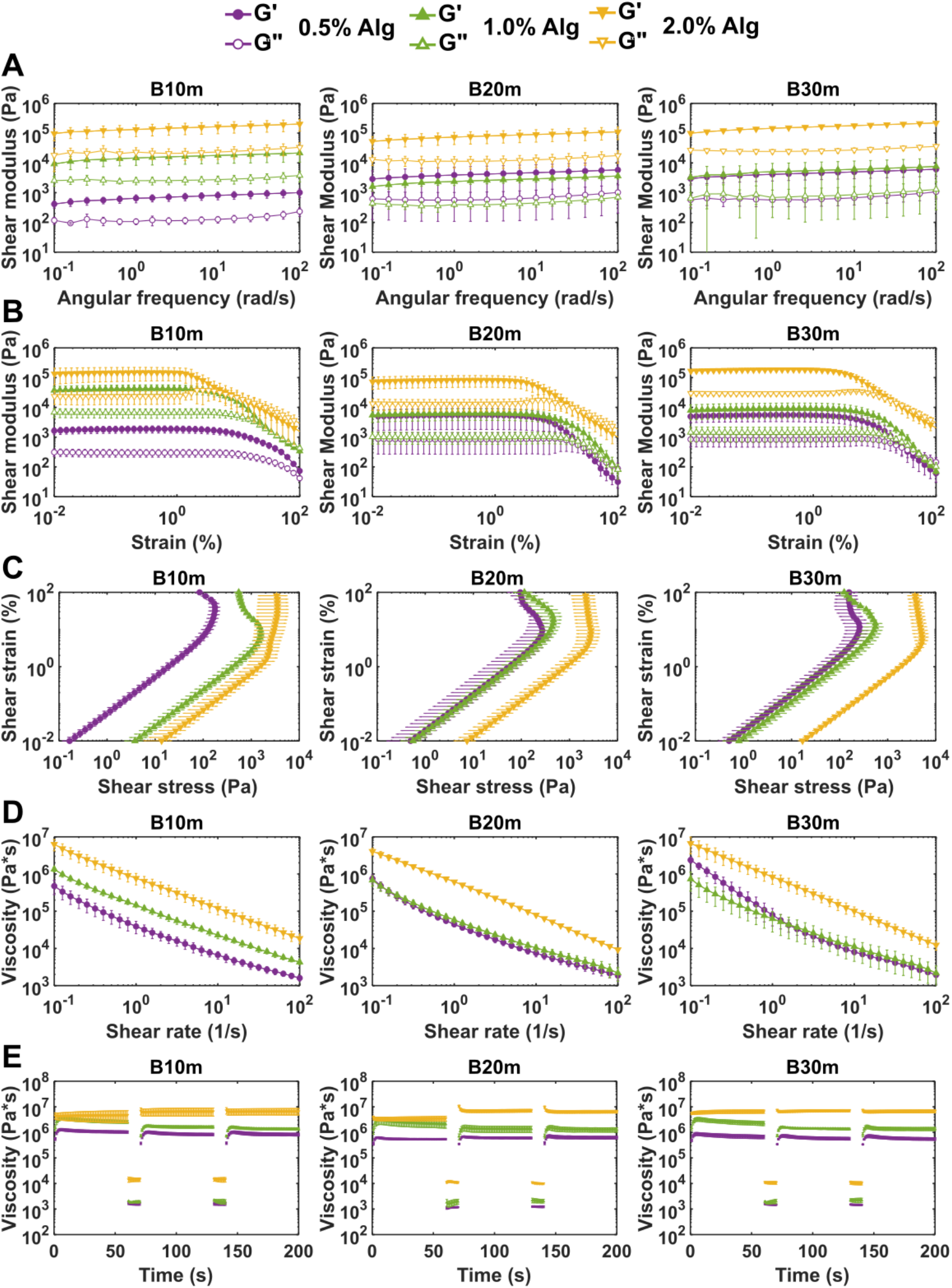
Rheological characterization of Alg microgels at different concentrations and blending times (*n* = 3). Measured shear modulus from (A) frequency sweep of microgels at an angular frequency ranging from 0.1 to 100 rad s^-1^; and (B) amplitude sweep of microgels at a strain ranging from 0.01 to 100 %; (C) Shear strain vs. Shear stress curves for Alg microgels; (D) flow curves of Alg microgels from rotational testing at a shear rate ranging from 0.1 to 100 s^-1^; (E) viscosity measurement at five intervals at alternating shear rates (the shear rate of 0.1 s^-1^ for 60 s and 100 s^-1^ for 10 seconds).

Dynamic viscoelastic properties from the frequency sweep test are presented in figure 3(A). All groups showed superior storage modulus G’ indicating elastic properties, with regard to loss modulus G”, (G’ > G”) over the entire angular frequency range 0.1-100 rad s^-1^. Besides, G’ was elevated when the concentration of Alg increased indicating that higher Alg concentrations possessed higher mechanical rigidity. The shear moduli of 2.0% Alg were considerably higher than those for 0.5 and 1.0% Alg.

The amplitude sweep test, as depicted in figures 3(B) and (C), illustrates the physical characteristics of Alg microgels. The results demonstrated flow points of Alg microgels - where the material transitioned from an elastic gel state (G’ > G”) to a viscous liquid state (G” > G’). Preceding the solution-gelation (sol-gel) transition point (at G’ = G”), all Alg microgels showed linear viscoelastic behavior. 2.0% Alg had a lower flow point indicating that 2.0% Alg was easily deformed at the same amount of strain compared to 0.5 and 1.0% Alg. However, as the blending time increased, the sol-gel transition points of 0.5 and 1.0% Alg were elevated to a similar level, which demonstrated that 0.5% Alg (B10m, B20m, and B30m), and 1.0%Alg (B20m and B30m) could not be easily deformed.

Using the amplitude sweep test, the shear strain – shear stress graphs were plotted to evaluate the yield stress of Alg microgels. In figure 3(C), the shear stress increased up to a certain point of shear strain and then decreased afterwards. All plots had this inflection point, indicating the yield stress point. For 2.0% of Alg with B10m, a continuous hardening stage was observed even beyond the yield-stress point. By increasing the blending time, 2.0% Alg experienced a softening stage. 2.0% Alg microgels had significantly higher yield stress regardless of the blending time compared to the yield stress of 0.5 and 1.0% Alg (Supplementary figure 1). 1.0% Alg with increased blending time showed a significant decrease in the yield stress from 993 to 223 (B20m) and 298 Pa (B30m). 0.5% Alg did not show major differences in the yield stress for different blending times compared to 1.0% Alg. 0.5% Alg (B10m, B20m, and B30m) and 1.0% Alg (B20m and B30m) had appropriate yield stress so that spheroids could not only be transferred in microgels without damage, but also to be maintained in their desired positions after bioprinting.

Viscosity versus shear rate (ranging from 0.1-100 s^-1^) plots were generated using the flow sweep test to investigate the shear-thinning behavior of Alg microgels (figure 3(D)). All Alg microgels possessed shear-thinning behavior, where the viscosity diminished with shear rate, as microgels were favorably rearranged along the flow direction (20). The viscosity of Alg was substantially augmented by increasing the Alg concentrations in the entire range of shear rate without sacrificing shear-thinning properties. When the blending time increased, the viscosity difference between 0.5 and 1.0% of Alg decreased. However, 2.0% Alg exhibited extremely high viscosity and shear modulus. Therefore, 2.0% Alg microgels with increased blending time did not exhibit significant changes in their viscosity as much as 0.5% Alg microgels under a high shear rate (100 s^-1^).

While bioprinting, the support bath was deformed due to the high shear rate during the needle movement. A support bath with self-healing characteristics quickly recovers from the damaged zone induced by the nozzle movement, making self-healing properties of the support gel essential for AAfB. Self-healing properties were examined through five cycles applied at two different alternating shear rates: low (0.1 s^-1^) – high (100 s^-1^) – low (0.1 s^-1^) – high (100 s^-1^) – low (0.1 s^-1^) as demonstrated in figure 3(E). The low shear rate (0.1 s^-1^) was applied for 60 s to imitate the stationary state of the support bath without the needle movement, whereas, the high shear rate (100 s^-1^) was applied for 10 s to mimic the state experienced during the needle movement. The self-recovery behavior was observed by the difference between before and after applying the shear rate. For 2.0% Alg, viscosity was elevated whenever a high shear rate was applied indicating shear-thickening behavior. When the support bath possessed shear-thickening behavior, the material alters its properties after the placement of spheroids. This poses limitations during repetitive bioprinting of spheroids, especially required for fabrication of scalable tissues. 1.0% Alg exhibited insufficient self-healing behavior. After the high shear rate mode, the viscosity did not fully recover. On the other hand, 0.5% Alg showed appealing recovery properties for all blending times. These rheological properties enabled spheroids to be bioprinted without inducing any major deformations for the gel and spheroids. Based on the assessed rheological properties, 0.5 and 1.0% Alg microgels with 10, 20, and 30 m blending time were selected for further consideration in spheroid bioprinting experiments.

### 3.3 Spheroid printability and fusion efficiency

For enabling successful self-assembly of bioprinted spheroids, positional accuracy and precision of spheroids is crucial. Figure 4(A) presents the positional accuracy and precision with respect to the size of bioprinted spheroids. The positional accuracy was improved with increased blending time such that 8.8, 5.4, and 5.2% positional accuracy was attained for 0.5% Alg with 10, 20, and 30 m blending time, respectively. 1.0% of Alg had a lower precision and accuracy compared to 0.5% Alg, where 16.8, 10.6, and 9.1% positional accuracy was achieved for 10, 20, and 30 m blending times, respectively. The positional precision for 0.5% Alg (B10m, B20m, and B30m) and 1.0% Alg (B10m, B20m, and B30m) was determined to be 7.4%, 6.6%, 4.7%, 20.4%, 12.2%, and 8.3%, respectively, where the precision increased with increasing blending time. The highest accuracy and precision was achieved with 0.5% Alg at B30m, thus, the smallest microgel particles yielded the highest positional accuracy and precision.

**Figure 4.**
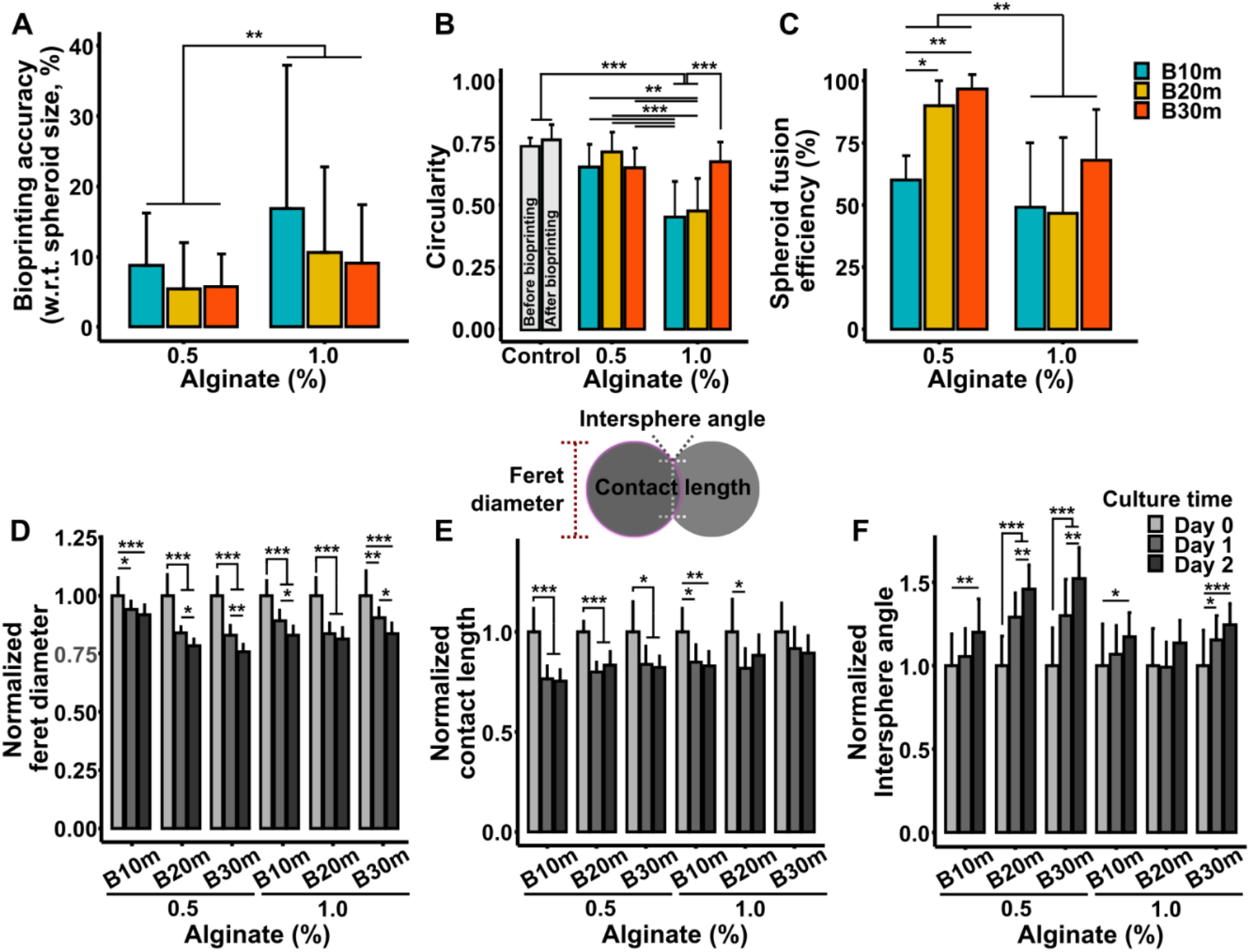
Characterization of bioprinted hMSC spheroids in Alg microgels. (A) Positional accuracy (mean) and precision (standard deviation) of bioprinted spheroids (*n* = 10, \**p* < 0.05, \*\**p* < 0.01, and \*\*\**p* < 0.001); (B) quantitative assessment of the deformation of bioprinted spheroids in Alg microgels (*n* = 9); (C) the fusion efficiency of hMSC spheroids (*n* = 10, \**p* < 0.05, \*\**p* < 0.01, and \*\*\**p* < 0.001); assessment of spheroid fusion dynamics for 2 days in Alg microgels: (D) ferret diameter, (E) contact length, and (F) intersphere angle (*n* = 10, \**p* < 0.05, \*\**p* < 0.01, and \*\*\**p* < 0.001).

The deformability of spheroids is yet another important factor that is critical for successful bioprinting (8). When the support bath had a higher mechanical strength, aspirated spheroids were not able to be bioprinted within the support bath owing to the high resistance of Alg microparticles, leading to low positional accuracy and precision. Otherwise, spheroids could be forcibly dragged to the desired locations in the stiff support bath with increased aspiration pressure, but this would induce structural deformation and decreased viability in spheroids. In this regard, we performed experiments to explore the role of concentration and blending time on the deformation of spheroids during bioprinting. In figure 4(B), the deformation of spheroids was represented by circularity. The circularity was quantitively analyzed with optical microscopic images of bioprinted spheroids (Supplementary figure 3). Before bioprinting, fabricated spheroids did not have a perfectly circular shape (circularity: 0.74 ± 0.03). As a secondary control group, spheroids were aspirated with 70 mmHg back pressure and placed back into the media yielding a circularity of 0.76 ± 0.06, which implies that 70-mmHg aspiration pressure did not significantly affect the morphology of spheroids. The circularity of bioprinted spheroids in 0.5% Alg with B10m, B20m, B30m, and 1.0% Alg with B30m were observed to be similar to that of the control groups, indicating preservation of circularity and retention of the spherical shape. In contrast, bioprinted spheroids in 1.0% Alg with 10 m and 20 m in blending time induced major damage to the spherical shape. In 1.0% Alg, when the blending time increased, the circularity of bioprinted spheroids was improved. The result implies that the smaller particles reduced the excess yield stress of 1.0% Alg microgels and augmented the rheological properties, allowing spheroids to be bioprinted without substantial deformation. In order to test spheroid fusion efficiency, dumbbell shapes were generated using hMSC spheroids as shown in figure 4(C) and figures 1(C1)-(C2). Here, the spheroid fusion efficiency implies the percentage of successfully fused spheroid pairs after a 2-day culture. The spheroid fusion efficiency for 0.5% Alg was ~60, 90, and 97% for B10m, B20m, and B30m, respectively. For 1.0% Alg, spheroids bioprinted in Alg microgels had ~49 (B10m), 47 (B20m), and 68% (B30m) fusion efficiency. With increased blending time, spheroid fusion efficiency was improved.

To characterize the dynamic fusion behavior in Alg microgels, ferret diameter and contact length of spheroids and intersphere angle between spheroids were analyzed (Supplementary figure 2 and Supplementary video 3). As demonstrated in figure 4(D), bioprinted spheroids in all Alg microgels exhibited shrinkage over 2 days, consistent with other studies (11). Considering that the diameter of spheroids became smaller one day post bioprinting, contact length also decreased (figure 4(E)). On Day 2, the contact length of assembled spheroids showed negligible difference compared to that on Day 1. Figure 4(F) demonstrates the intersphere angle between fusing spheroids. Regardless of the diameter and contact length, the intersphere angle increased. Notably, bioprinted spheroids in 0.5% Alg with B30m showed dramatically increased intersphere angle indicating that bioprinted spheroids fused well in this group. In 1.0% Alg microgels with B10m and B20m, the intersphere angle did not change over two days, probably due to the rigid properties of 1.0% Alg microgels. However, the intersphere angle in 1.0% Alg microgels increased when the blending time increased.

As presented above, 0.5% Alg with 30 m blending time exhibited favorable rheological characteristics for AAfB enabling highest positional precision and accuracy, and spheroid fusion efficiency without inducing major deformation on the morphology of spheroids during bioprinting. Thus, we preferred to use 0.5% Alg with 30 m blending time for the rest of the experiments.

### 3.4 AAfB of hMSC spheroids for bone tissue fabrication

With the optimized Alg microgels (0.5% Alg with B30m), we bioprinted hMSC spheroids and osteogenically inducted them to demonstrate bone tissue formation. To systematically study the influence of the addition of the osteoclastogenically inducted monocytes, we also bioprinted the 2:1 hMSCs:THP1 and 3:2 hMSCs:THP1 in a similar fashion. These spheroids were cultured in their respective ration of differentiation media for 24 h and then bioprinted into a triangular shape as discussed before. After bioprinting, all the structures were cultured up to 28 days. H&E and alizarin S red staining and immunostaining (figures 5(A) and (B)) with osteoblast and osteoclast specific markers proved that both hMSC only and THP1 involved structures showed compact spheroid arrangements. The hMSC only spheroids were observed to compact significantly and form a tissue ball (figure 5(A)) over the 28-day culture period, while the 2:1 hMSCs:THP1 spheroids were observed to maintain the desired triangular shape during the entire course of culture. All groups were positive for calcium deposition (from alizarin S red staining), with higher intensity observed in the 2:1 hMSCs:THP1 group compared to the hMSC only and 3:2 hMSCs:THP1 groups (figure 5(B)). Immunofluorescent staining showed that both RUNX2 and TRAP staining were positive in the heterocellular spheroids indicating osteogenic and osteoclastogenic differentiation, respectively, with homogeneous distribution of both cell types inside the spheroids. Osteoclastogenically-inducted cells were located more in the periphery than the core of the structures in both 2:1 hMSCs:THP1 and 3:2 hMSCs:THP1 groups. qRT-PCR study shows expression of both osteoblast and osteoclast specific bone markers, indication appropriate differentiation of hMSCs and THP1 into osteoblasts and osteoclasts, respectively. 3:2 hMSCs:THP1 group exhibited increased expression levels for the BSP gene (~6745-folds) and MMP9 gene (~24662-folds) at Day 28 as compared to those in the hMSC only group. The expression level of the RUNX2 gene (8-folds) for the 2:1 hMSCs:THP1 group was significantly higher than that of the hMSC only group. Overall, most of the osteogenic and osteoclastogenic genes in 2:1 hMSCs:THP1 and 3:2 hMSCs:THP1 groups showed greater level of expression as compared to those in the hMSC only group.

**Figure 5.**
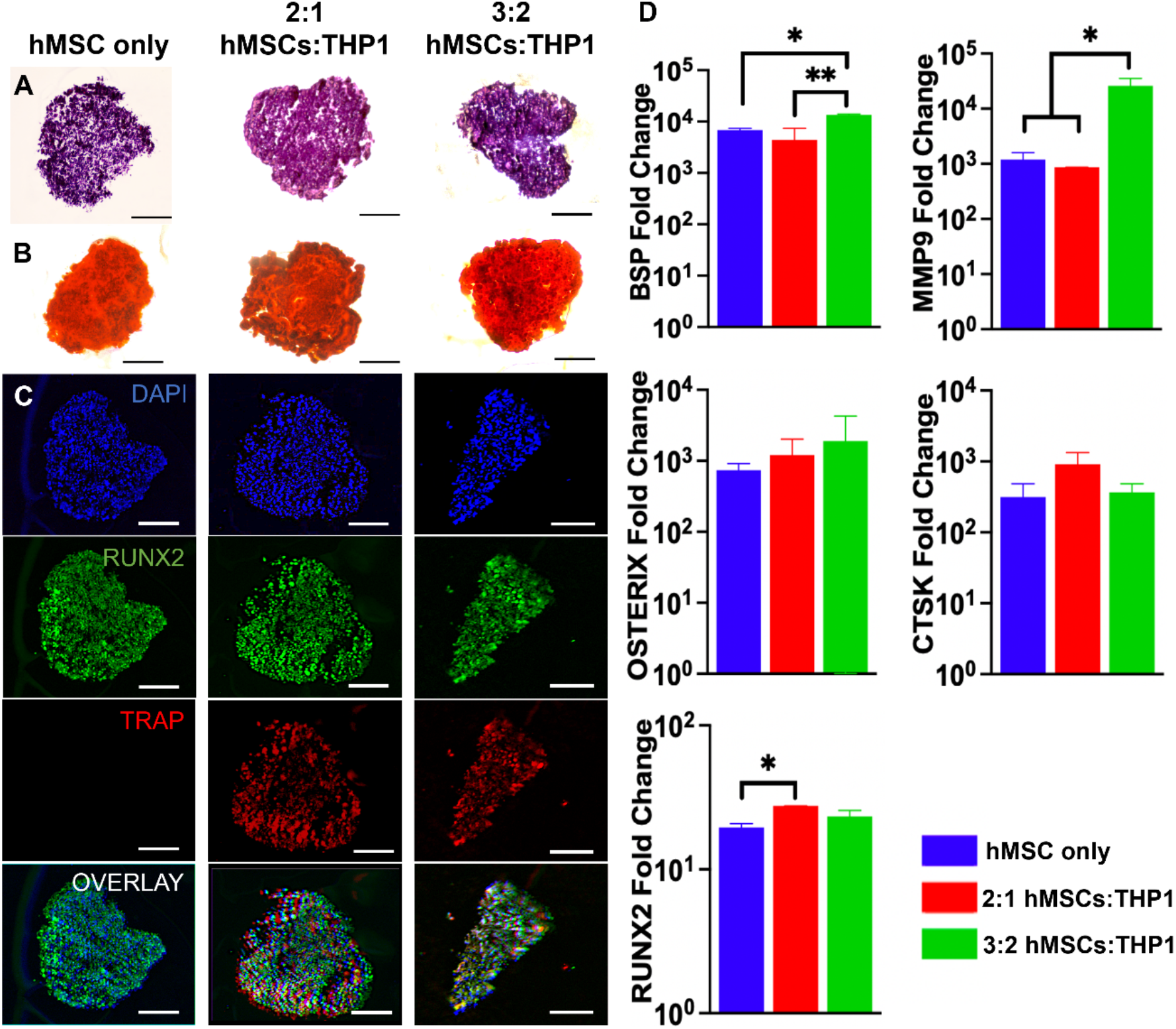
Bioprinted stem cell spheroids for bone tissue fabrication. (A) H&E and (B) alizarin S red staining images; (C) Immunostaining with RUNX2 for hMSCs differentiating into osteoblasts and TRAP for THP1 differentiating into osteoclasts; (D) Quantification of BSP, OSTERIX, RUNX2, MMP9, and CTSK gene expressions of bioprinted bone tissues (*n* = 3, \**p* < 0.05 and \*\**p* < 0.01). Scale bar: 200 μm.

## 4. Discussion

In the field of tissue engineering and regenerative medicine, artificial functional scaffolds with complex geometry and orientations are often required to mimic native tissues. To achieve this goal, freeform bioprinting with supporting materials has been explored as an option (5, 9, 11–13, 21–24). Recently, our group has introduced AAfB as a promising bioprinting technique to fabricate tissue structures with complex 3D patterns of spheroids (8, 17). However, an ideal support bath for enabling adequate spheroid fusion and transparency has still been a challenge. For developing a support bath, an appropriate understanding of the rheological properties is essential, and without such exploration, it is difficult to predict whether the material has adequate yield stress to support bioprinted structures. Moreover, it could not also be confirmed that the microgels have sufficient self-healing property which is crucial for allowing the recovery of the damaged region after bioprinting. Inferior self-healing behavior induces major deformation of the shape of microgels, especially during scalable tissue fabrication when there is a significant back-and-forth motion of the needle. Therefore, we systematically investigated the rheological properties for the development of a supporting bath with Alg microgels to demonstrate enhanced functionality for the bioprinting of spheroids without adversely affecting their viability and shapes, resulting from high shear stress during the AAfB process.

The proposed supporting bath is composed of microgels. The microgel-microgel interactions are reduced when the shear rate is increased such as moving a needle inside the bath. Therefore, as observed in figures 3(D) and (E), the viscosity decreased when higher shear rates were applied. However, Alg microgels were able to rearrange themselves after exposure to higher shear rates (when the needle was moving inside the bath) by efficiently packing and forming the originally organized shapes due to self-healing characteristics. The shear-thinning yield-stress gel immobilized the spheroid in place and showed remarkable flowability when the needle was moving inside the bath. Supplementary figure 4 demonstrates how Alg microgels deformed directly after the bioprinting of spheroids. The scattered light on the uneven surface caused by the deformed microgels appeared in dark color. For 1.0% Alg, the black region in the images indicated the damaged microgels by the needle during bioprinting due to the low self-healing behavior (figure 3(E)). The 1% Alg with 10 m blending time group exhibited a high yield stress owing to the larger particle size. However, the increased blending time reduced the yield stress (Supplementary figure 1). The yield stress also influenced the bioprinting precision and accuracy, and the immoderate yield stress induced a morphological change on bioprinted spheroids (figure 4(B) and Supplementary figure 3). For 0.5% Alg, however, even after bioprinting, Alg microgels enabled neatly positioning of spheroids without any permanent deformation on the gel. In contrast, with 2.0% Alg, spheroids could not be easily transferred into the support bath (data not shown) due to its high yield stress (figure 3). Even if the spheroids were able to penetrate into the gel, bioprinted spheroids lost their original shape. In rheological assessments, the viscosity of 2.0% Alg was elevated whenever a high shear rate was applied (figure 3(E)). The amplitude sweep test (figure 3(B)) demonstrated that 2.0% Alg microgels were easy to deform even though a low strain level was applied indicating that 2.0% Alg microgels possessed a shear-thickening behavior, which is not suitable for AAfB. In this regard, 2.0% Alg did not exhibit a strain-softening in the shear stress – shear strain curve (figure 3(C)). Therefore, 2.0% Alg was not considered for spheroid bioprinting. In figure 3(D), the initial viscosity (at 0.1 s^-1^) of 0.5% Alg was increased with increasing blending time. The smaller size particles yielded higher viscosity due to the increased probability of particle-to-particle interactions (25). However, 1.0% and 2.0% Alg microgels might exhibit not linear rheological properties due to the viscosity of the material itself and their larger particle size compared to 0.5% Alg microgels (26). In morphological analysis, Alg microgels were not uniform but polydispersed (figures 2(A) and (B)). Therefore, the trend of the rheological behavior was not expected to be linear in polydisperse microgels. The lower shear modulus of 0.5% Alg facilitated the blended Alg microgels to generate a narrow particle size distribution (figure 2(B)). Even though the rheological analysis allowed us to obtain extensive data, profound and sophisticated analysis of spheroid motion is also required to reveal the interactions between microgels and the nozzle with an aspirated spheroid during bioprinting. Moreover, there is always a possibility of differences between the rheological assessment and actual bioprinting results. In the actual bioprinting process, additional factors, such as gravity and surface tension driven forces between air and Alg microgels might play a role.

Alginate is a natural substance extracted from brown algae and is widely used in various fields as a cost-effective, highly biocompatible, and easily handled biomaterial. Because of its known excellent biocompatibility (4), hMSC spheroids were viable at all time points over the three day culture and the presence of dead cells in all groups was negligible (Supplementary figure 6). The reduced sizes of Alg microgels with increased blending time did not affect the cell viability. For extrusion-based bioprinting (EBB), 2-4% of alginate has been commonly used owing to adequate mechanical stability post bioprinting (4). In such cases for EBB, alginate is crosslinked with a lower concentration of calcium ions (depending on the crosslinking agent, e.g. 0.1-0.5% CaCl_2_ (27), 1.0% CaSO_4_ (28), or 0.2% CaCO_3_ (24)) to maintain the printed shape after extrusion, followed by further addition of a higher content of calcium ions to crosslink the bioprinted structures. However, for AAfB, a lower concentration (< 1.0%) of alginate is desired not only to protect spheroids during bioprinting but also to obtain optimal mechanical and rheological properties suitable for its use as a support material. Apart from the crosslinking strategy for EBB, fully-crosslinked alginate gels are also utilized as a support material by blending them for optimized time periods (9, 11). Therefore, a proper combination of blending time and concentration of alginate is crucial to modulate the microgel properties for AAfB.

In our recently reported work (11), 0.5% Alg microgels with 10 m blending yielded a 35% spheroid positional accuracy with respect to the spheroid size; however, for this study, owing to changes in preparation protocol, the same combination yielded an accuracy of ~9%. Even though we observed a higher positional accuracy, Alg microgels disturbed the spheroid fusion by getting entrapped between bioprinted spheroids. Thus, the efficiency of spheroid fusion was < 65% with 0.5% Alg with B30m. However, we here demonstrated more than 90% fusion efficiency with decreased particle size. With increased blending time, Alg microgels with considerably improved yield stress, shear thinning, and self-healing properties were attained, with higher overall bioprinting ability with no negative effects on cellular activities. In the previous work (11), prepared alginate microgels were not transparent, making bioprinting or further imaging process extremely challenging. In this study, the modified preparation procedure for Alg microgels provided us with the desirable transparency of Alg microgels, adequate for AAfB. Although we here demonstrated Alg microgels with optimal properties for spheroid bioprinting, a commercial blender was used to minimize Alg microgel size. However, this process yielded highly irregular shapes. Therefore, it will be ideal to use other methods (such as microfluidics) to produce uniform micron-size particles without blending, which will be considered for our future studies.

A detailed understanding of the underlying physics and a simulation study for AAfB with Alg microgels based on the rheological properties will be highly beneficial as the bioprinting parameters, such as bioprinting speed and aspiration pressure, are dependent on the rheological properties of the microgels. When the support material has excessive yield stress and viscoelasticity, biological activities of bioprinted spheroids deteriorate, even though the spheroids can be successfully positioned in the gel. In addition, an optimal aspiration pressure for spheroid picking is important. The excess back pressure is often associated with the deformation of the spheroids during bioprinting. As observed in Supplementary figure 3, bioprinting with. an aspiration pressure of 70 mmHg and bioprinting speed of 2.5 mm s^-1^ in 0.5% Alg microgels did not induce any substantial change in the spheroid shape and viability; thus, these bioprinting parameters were fixed throughout the entire study. With the optimized rheological properties and morphological assessment of blended Alg microgels with different concentrations of alginate, multiple spheroids could be easily bioprinted with over >90% spheroid fusion efficiency. It is worth mentioning that the initial fusion strength of the spheroids is low right after bioprinting, and high pipetting pressure might break the fused spheroids while transferring them into the new culture flasks or during media change. Thus, fused spheroids should be treated gently in culture.

AAfB of spheroids has been utilized to generate triangular patterns using hMSC and heterocellular hMSCs:THP1 spheroids as a proof of concept study using Alg microgels to demonstrate fabrication of tissue patches and spheroid fusion efficacy. During the culture of bioprinted hMSC spheroids, the spheroid size slightly decreased as shown in figure 4(D), Supplementary figure 2, and Supplementary video 3. This is because bioprinted spheroids in Alg microgels were inclined to gain compactness without proliferation and migration into Alg microgels due to the lack of cell-binding peptides of Alg, which is in line with a previous study (29). In 1.0% Alg microgels with B10m and B20m, the intersphere angle was not extensively changed over two days, probably due to the rigid properties of 1.0% Alg microgels.

hMSC spheroids are reported to readily self-assemble forming tissue patches (11, 30). The degree of spheroid fusion can be modulated by optimizing the differentiation time period. The spheroids are prone to maintain their shape more upon differentiation towards osteogenic lineage, with a decreased fusion efficacy. To build up a large-scale structure, it is essential for bioprinted spheroids to fuse to each other; otherwise, they can break apart. Therefore, in this work, we studied the optimization of Alg microgels for increasing the integrity of bioprinted spheroids and in turn, improved the fusion efficiency. Depending on the fusion from Day 1 to 5, Alg microgels were gently removed by immersing in sodium citrate as an alginate lyase (Supplementary figure 5) followed by washing with DPBS after fusion. The spheroids were then cultured in their individual differentiation media for 28 days to allow differentiation of hMSCs and THP1 into osteoblasts and osteoclasts, respectively. Figures 5(A) and (B) presented fused bioprinted tissue patches fabricated with hMSC only and 2:1 and 3:2 hMSCs:THP1 heterocellular spheroids. H&E images demonstrated compact structure for all groups. THP1-involved groups were observed to demonstrate more retention of the bioprinted shape compared to the hMSC only group. The triangular shape was clearly visible in 2:1 hMSCs:THP1 group. The hMSC only group formed a tissue ball over the 28-day culture period due to more compaction of hMSC spheroids. Calcium deposition was noticed in all the groups from positive alizarin S red staining. Notably, more calcium deposition was observed in 2:1 hMSCs:THP1 group compared to the other groups indicating more bone tissue formation. Both TRAP and RUNX2 staining were positive for the hMSCs:THP1 spheroids, with uniform distribution of RUNX2 across the entire bioprinted structure domain, while the TRAP staining was more concentrated towards the periphery of structures. This is probably owing towards more compaction potential of the hMSCs, enabling them to occupy the core of the spheroids, whilst the THP1 lined up the periphery. In fact, THP1 alone did not have the potency to self-assemble and form spheroids over a period of 48 h (data not shown). Immunostaining also demonstrated stronger expression of RUNX2 and TRAP activity for the 2:1 hMSCs:THP1 spheroids compared to the other groups. Thus, we showed that the bioprinted structures with heterocellular spheroids exhibited both osteogenic and osteoclastogenic properties, with more calcium deposition and shape retention for the 2:1 hMSCs:THP1 group. Expression of osteogenic and osteoclastogenic genes were investigated for different groups of spheroids, including BSP, OSTERIX, RUNX2 and MMP9 and CTSK at Day 28. All the groups exhibit expression of osteogenic and osteoclastogenic markers, corroborating the immunostaining data. Compared to the hMSC only group, the 2:1 hMSCs:THP1 group showed higher expression of the late- and mid-stage osteogenic markers - RUNX2 and OSTERIX respectively, with a lower expression of the early stage marker BSP, indicating a higher degree of hMSC differentiation into osteoblasts. Significantly higher expression of MMP9 in 3:2 hMSCs:THP1 group, on the other hand, indicates higher degree of extracellular matrix degradation (31), which may not be ideal for the fabrication of bone-tissue patches. Thus our results indicate that bone tissue fabricated by AAfB of 2:1 hMSCs:THP1 spheroids provided a more biomimetic scenario to study bone remodeling *in vitro*.

The presented study aimed to improve the AAfB process for tissue spheroid placement in 3D and determine the optimal rheological properties to obtain an appropriate support bath for AAfB. As an application, we also presented the fabrication of bone tissue patches using heterocellular hMSCs:THP1 spheroids expressing osteogenic and osteoclastogenic genes as building blocks. Using these optimized Alg microgels, various types of spheroids can be used for the fabrication of scalable and highly complex geometries of tissues and organs.

## 5. Conclusion

In this study, Alg microgels have been thoroughly investigated to support aspiration-assisted freeform bioprinting in a biocompatible and transparent medium, where decreased microgel size facilitated higher bioprinting accuracy and precision as well as spheroid fusion efficiency without inducing major deformation and cell death during bioprinting. Heterocellular stem cell spheroids were bioprinted in the proposed Alg microgels, where bioprinted structures exhibited successful self-assembly of spheroids with shape retention, and expression of both osteogenic and osteoclastogenic genes. Thus, our results indicate that the proposed Alg microgels have the potential to become a prospective support material for freeform bioprinting of tissue spheroids with remarkable spheroid fusion efficiency and bioprintability, demonstrated here with bone tissue fabrication as a proof of concept.

## Acknowledgements

This work has been supported by National Science Foundation Award 1914885 (I.T.O.), National Institute of Dental and Craniofacial Research Award R01DE028614 (I.T.O.), and a contract from The Lysosomal and Rare Disorders Research and Treatment Center Inc. (I.T.O.). We thank RoosterBio for providing hMSCs and growth media, and Bio-Techne for providing several reagents in support of our experiments.

