## Supplementary Materials for "Aspiration-assisted freeform bioprinting of mesenchymal stem cell spheroids within alginate microgels"

**Supplementary Videos**

**Supplementary Video 1.** AAfB of hMSCs spheroids into dumbbell shape structures.

**Supplementary Video 2.** AAfB of hMSCs spheroids into pyramid shape structures.

**Supplementary Video 3.** hMSC spheroid fusion dynamics in dumbbell shape for six hours in 0.5% Alg with B30m.

**Supplementary Figures**

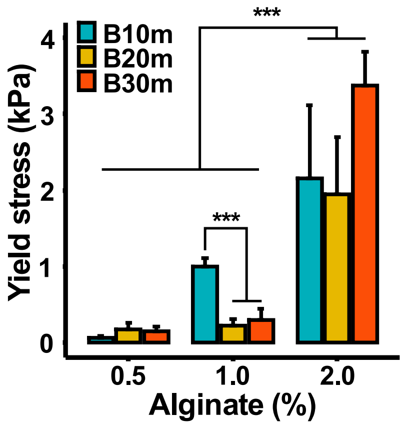


**Supplementary Figure 1.** Yield stress of Alg microgels (*n*=3; ***p<0.001).

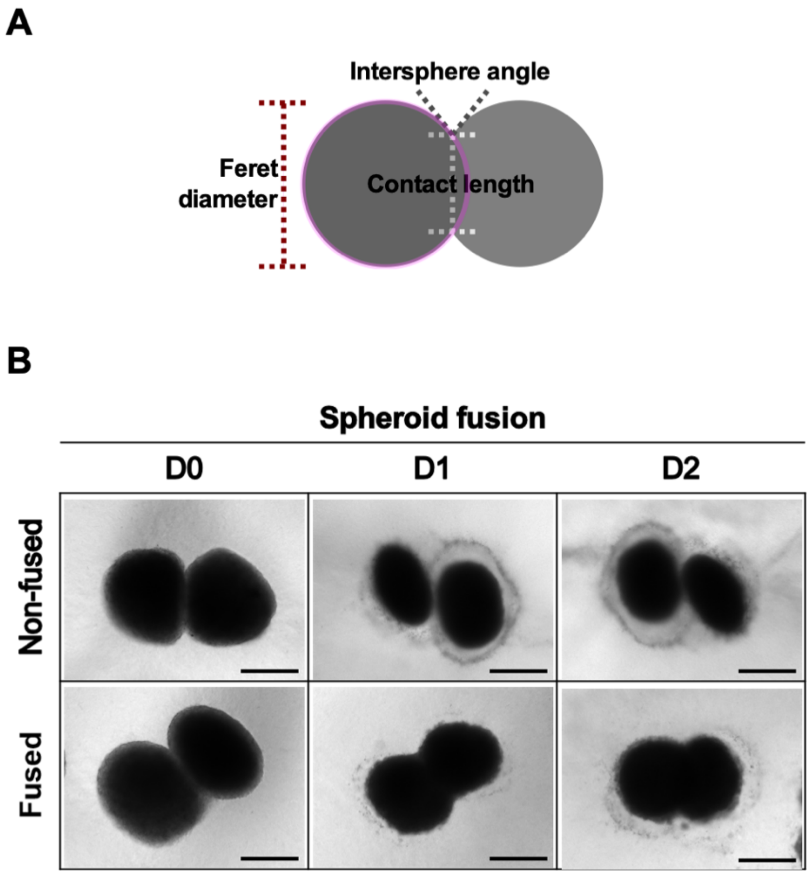


**Supplementary Figure 2.** (A) A schematic for quantitative assessment of spheroid fusion dynamics including diameter, intersphere angle, and contact length between bioprinted spheroids. (B) Demonstration of images of fused and non-fused spheroids during a two-day culture (D0: Day 1, D1: Day 1, and D2: Day 2). Scale bar, 500 µm.

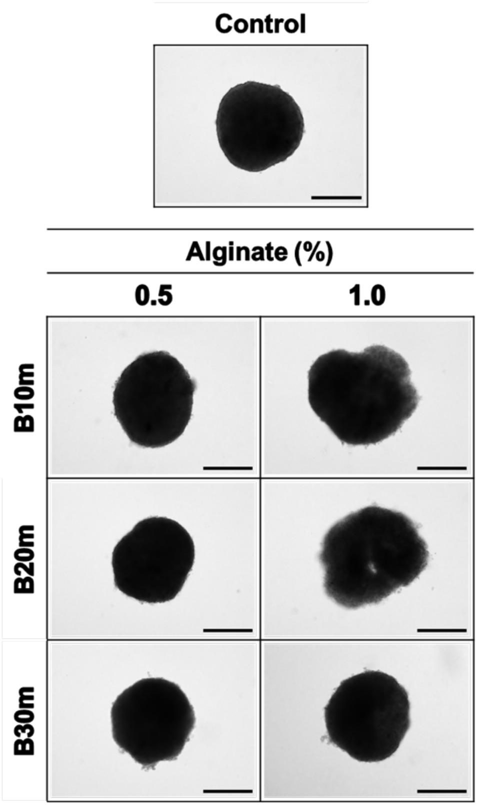


**Supplementary Figure 3.** Morphological shape of bioprinted spheroids in Alg microgels. Scale bar, 500 μm

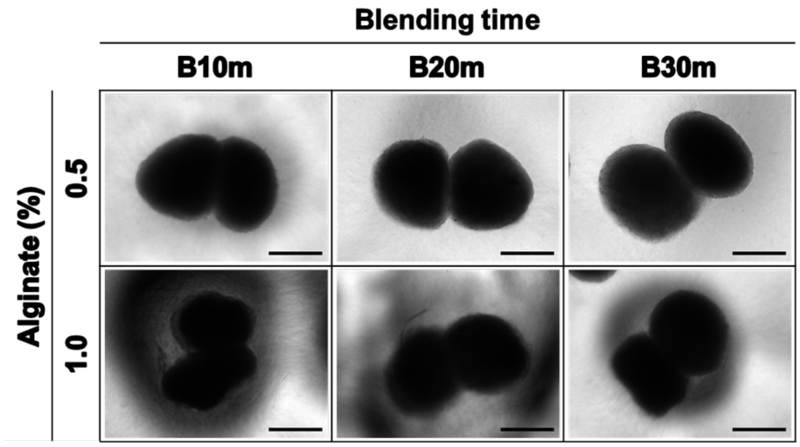


**Supplementary Figure 4.** Microscopic images of bioprinted spheroids in Alg microgels post bioprinting (Day 0). Black color in 1.0% Alg demonstrates deformed Alg microgels after bioprinting due to insufficient self-healing properties. Scale bar, 500 µm.

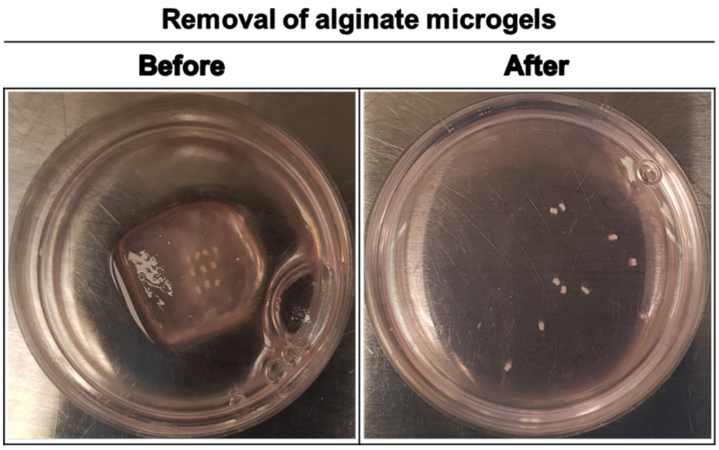


**Supplementary Figure 5.** The removal of Alg microgels (0.5% Alg with B30m) with the addition of 4% sodium citrate onto a few pairs of fully-fused dumbbell-shape structures.

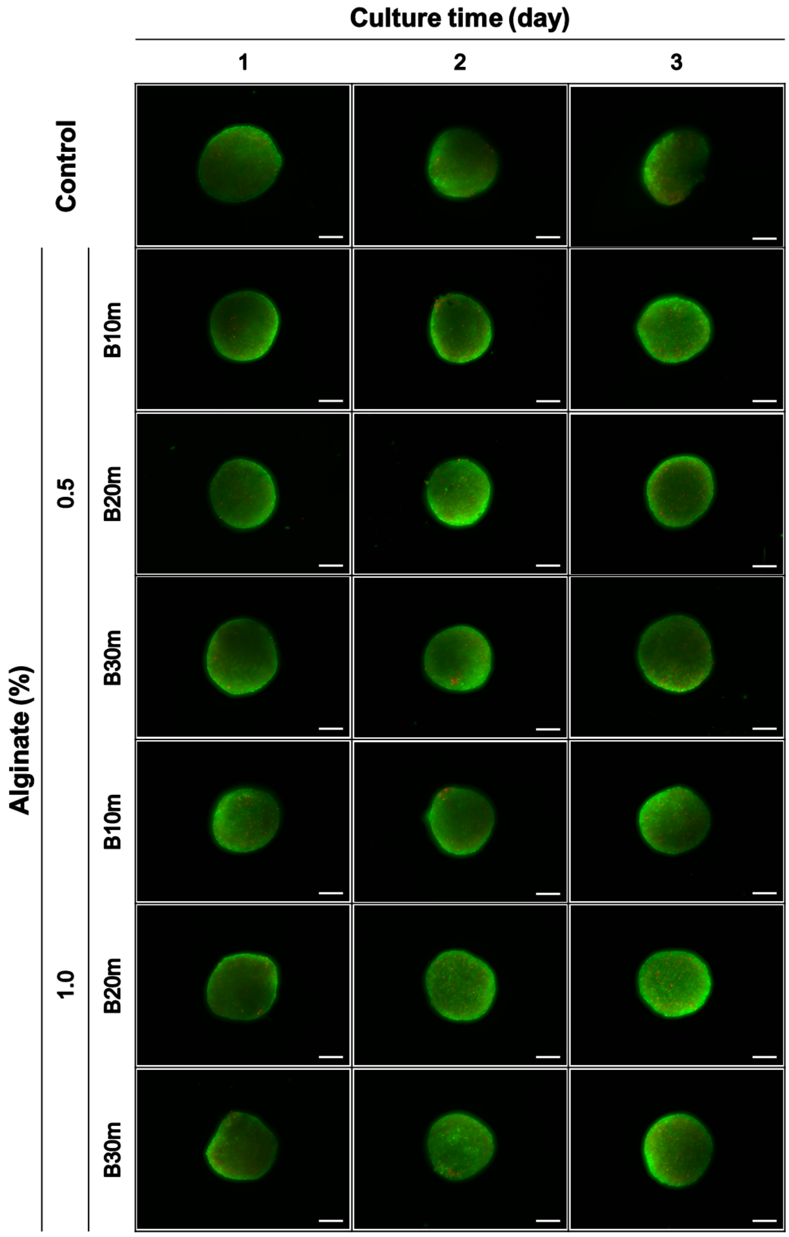


**Supplementary Figure 6.** Fluorescence images showing the biocompatibility of hMSCs spheroids cultured in Alg microgels during a 3-day incubation by LIVE/DEAD staining. Scale bars, 200 μm.
